# Robust and durable prophylactic protection conferred by RNA interference in preclinical models of SARS-CoV-2

**DOI:** 10.1101/2022.03.20.485044

**Authors:** Yesseinia I. Anglero-Rodriguez, Florian A. Lempp, James McIninch, Mark K. Schlegel, Christopher R. Brown, Donald J. Foster, Adam B. Castoreno, Tuyen Nguyen, Megha Subramanian, Martin Montiel-Ruiz, Hannah Kaiser, Anna Sahakyan, Roberto Spreafico, Svetlana Shulga Morskaya, Joseph D. Barry, Daniel Berman, Stephanie Lefebvre, Anne Kasper, Timothy Racie, Diann Weddle, Melissa Mobley, Arlin Rogers, Joseph Dybowski, Saeho Chong, Jayaprakash Nair, Amy Simon, Kevin Sloan, Seungmin Hwang, Herbert W. Virgin, Kevin Fitzgerald, Martin A. Maier, Gregory Hinkle, Christy M. Hebner, Akin Akinc, Vasant Jadhav

## Abstract

RNA interference is a natural antiviral mechanism that could be harnessed to combat SARS-CoV-2 infection by targeting and destroying the viral genome. We screened lipophilic small-interfering RNA (siRNA) conjugates targeting highly conserved regions of the SARS-CoV-2 genome and identified leads targeting outside of the spike-encoding region capable of achieving ≥3-log viral reduction. Serial passaging studies demonstrated that a two-siRNA combination prevented development of resistance compared to a single-siRNA approach. A two-siRNA combination delivered intranasally protected Syrian hamsters from weight loss and lung pathology by viral infection upon prophylactic administration but not following onset of infection. Together, the data support potential utility of RNAi as a prophylactic approach to limit SARS-CoV-2 infection that may help combat emergent variants, complement existing interventions, or protect populations where vaccines are less effective. Most importantly, this strategy has implications for developing medicines that may be valuable in protecting against future coronavirus pandemics.

---

The development of SARS-CoV-2 vaccines has been an extraordinary advance in the fight against the coronavirus disease (COVID-19) pandemic (*1–3*). However, the challenges behind distribution, adherence, global supply, and the rapid spread of the virus have led to the emergence of new variants and unremitting waves of infections globally. Moreover, immune response to vaccination may be highly variable in immunocompromised patients requiring orthogonal approaches to confer protection (*4*). Taken together, there remains an urgent need for the development of new antiviral tools to control viral spread and augment vaccination efforts.

RNA interference (RNAi) is a highly conserved natural mechanism utilizing siRNAs with cellular machinery to catalyze the sequence-specific downregulation of target messenger RNA (mRNA) or viral RNA (vRNA). After nearly two decades of effort, RNAi-based therapeutics have emerged as a new class of medicines with four approved drugs and additional agents in various stages of clinical investigation (*5–7*). In addition, recent advances in lung delivery have demonstrated that conjugation of 2’-*O*-palmityl (C16) to siRNAs delivered intranasally (IN) allows bronchiolar and alveolar uptake and durable silencing in the lung (*8*). The opportunity to target highly conserved regions of the SARS-CoV-2 genome with siRNA at the primary site of viral replication enables a potential broad and direct antiviral strategy. More importantly, by targeting the regions outside the spike gene, this approach could remain effective against emerging variants that often include mutations in the spike region. In addition, an RNAi approach may be efficacious in immunocompromised patients where vaccine response can be blunted. Here, we show that C16-siRNAs show protection against SARS-CoV-2 infection in vitro and in a hamster model when delivered prior to the infection.

## Results

### Design and screening of siRNAs

We assessed conserved sequences across SARS-CoV, SARS-CoV-2, and Middle East Respiratory Syndrome (MERS-CoV) genomes available in early 2020 and identified 1,511 fully conserved sites between SARS-CoV-2 and SARS-CoV genomes as well as 9 conserved sites between SARS-CoV-2 and MERS-CoV genomes (**Fig. S1A–B)**. A subset of 349 unique target sequences predicted to be amenable for RNAi was selected and the corresponding siRNAs were synthesized as C16-conjugates for further evaluation **(Table S1)**. Initial in vitro activity via transfection was assessed using two dual-luciferase reporter vectors where the (+) strand RNA SARS-CoV-2 target sites in a concatemerized arrangement were inserted into the 3’ UTR of *Renilla* luciferase (rLuc) while the firefly luciferase (fLuc) served as a control. In addition, a small subset of siRNAs targeting the (-) strand RNA at the open-reading frame 1 ab (ORF1ab) were also evaluated. This reporter assay identified 91 siRNAs with ≥80% target reduction relative to mock-transfected controls (**Fig. 1A)**. The siRNA sites with the highest activity were spread across the viral genome.

**Fig. 1.**
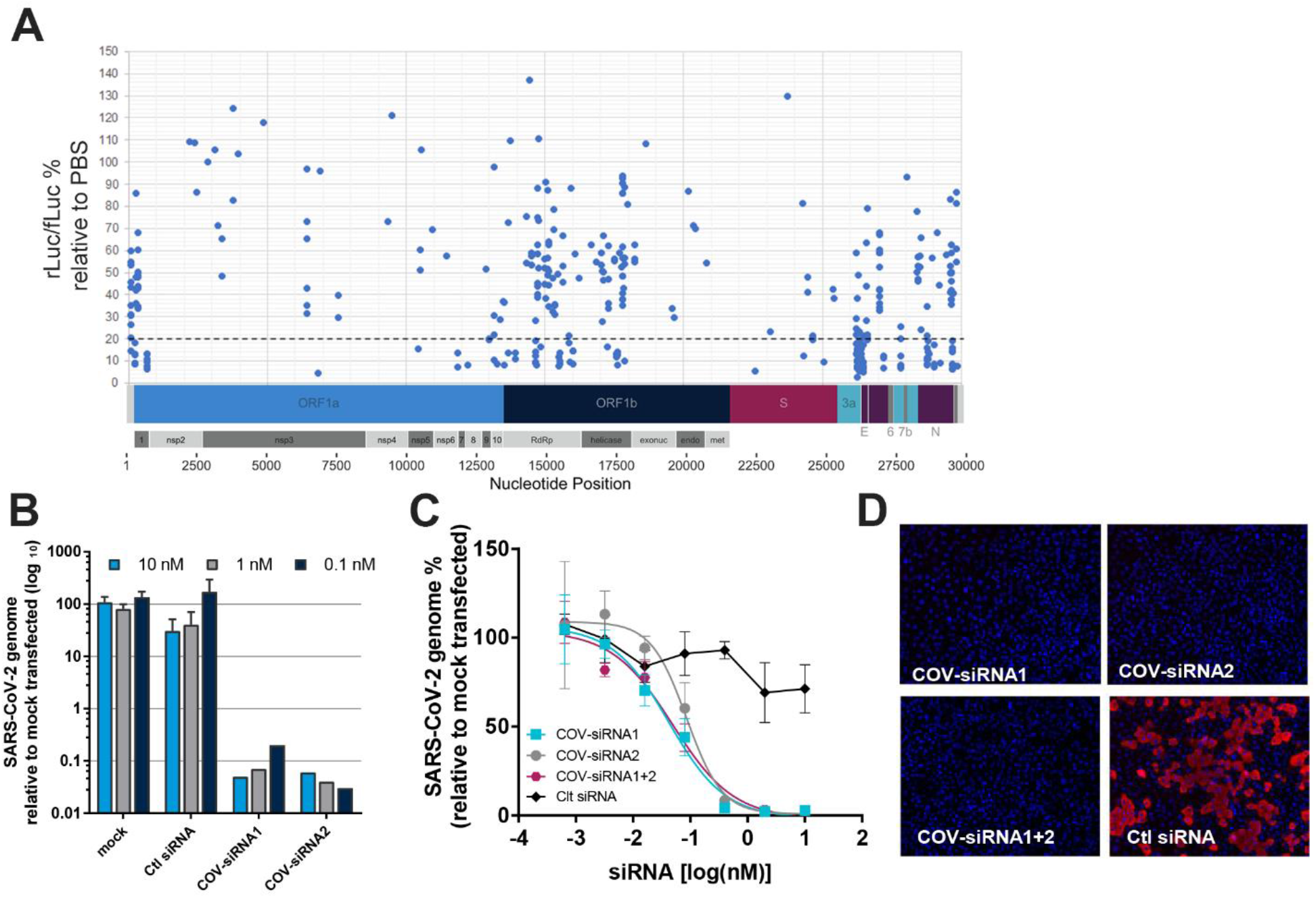
In vitro screening of SARS-CoV-2 targeted siRNAs. **(A)** Each dot represents the mean luciferase activity (rLuc/fLuc) relative to PBS in Cos7 cells from a single experiment with three independent repeats. Below the dotted line are siRNAs with ≥ 80% relative reporter repression. The diagram below represents the 30 kb SARS-CoV-2 viral genome and the target site region per siRNA. **(B)** For all infection assays, Vero E6 cells were reverse transfected with siRNA or controls and infected with SARS-CoV-2 isolate USA-WA1/2020. Twenty-four hours post transfection, cells infected, and supernatant was evaluated by RT-qPCR for viral genomic RNA from single experiment representative of one-four independent repeats ± SD. **(C)** Dose-dependent knockdown of viral genomic RNA after siRNA treatment quantified by RT-qPCR from two independent experiment with three independent repeats ± SEM. Cross-validated by quantitative IFA **(Fig. S2C–D)**. **(D)** Immunofluorescence at 10 nM concentration. SARS-CoV-2 N protein (red) and nuclei (blue).

The best performing siRNAs were evaluated in an in vitro infection assay using a live SARS-CoV-2. Vero E6 cells were transfected with siRNA prior to infection and virus in the supernatant was quantified by RT-qPCR. Twenty-three siRNAs demonstrated ≥2-log reduction in viral RNA at 10 nM, and 4 siRNAs demonstrated ≥3-log reduction in viral RNA at the lowest concentration tested, 0.1nM **(Fig. S2B; Fig. 1B).** Contrarily, none of the siRNAs targeting the (-) RNA genome showed activity in the live virus assay (**Fig. S2A**), consistent with the seemingly closed structure of the SARS-CoV-2 replication complex where the negative strand exists.

The top two siRNA candidates targeting within ORF1ab, were selected for further assessment, and demonstrated potent, dose-dependent antiviral activity, with EC_50_ values of 42 pM and 86 pM for COV-siRNA1 and COV-siRNA2, respectively **(Fig. 1C; Table S2)**. The activity was confirmed using a SARS-CoV-2 N protein immunofluorescence (IFA) assay **(Fig. S2C–D).** No intracellular viral N protein was detected in cells treated with COV-siRNA1, COV-siRNA2, or a combination (COV-siRNA1+2) at 10 nM **(Fig. 1D).**

### Two-siRNA combination prevents the emergence of SARS-CoV-2 escape mutants

The barrier to drug resistance was determined for the lead siRNAs using in vitro viral serial passaging methods (*9*, *10*). SARS-CoV-2 virus was passaged for five times in the presence of single-siRNA (COV-siRNA1) or the two-siRNA combination (COV-siRNA1+2), at 5x, 10x, and 20x EC_50_. The EC_50_ determinations were performed on pooled virus from each passage and compared to mock-treated control **(Fig. 2A–B**). With a single-siRNA treatment, viral escape mutants were evident at 5x EC_50_ with marked EC_50_ curve shifts by P4-P5. This trend was sustained at higher 10x and 20x concentrations, where viral escape mutants were observed at P3 and P2, respectively. In stark contrast, treatment with a two-siRNA combination continued to retain activity after serial passaging, suggesting a much higher barrier to escape. Even at P5, collected virus remained susceptible to COV-siRNA1 treatment comparable to virus serially passed with control siRNA **(Fig. S3)**. Overall, these data show that a two-siRNA combination targeting two distinct conserved sites created a formidable barrier to viral escape.

**Fig. 2.**
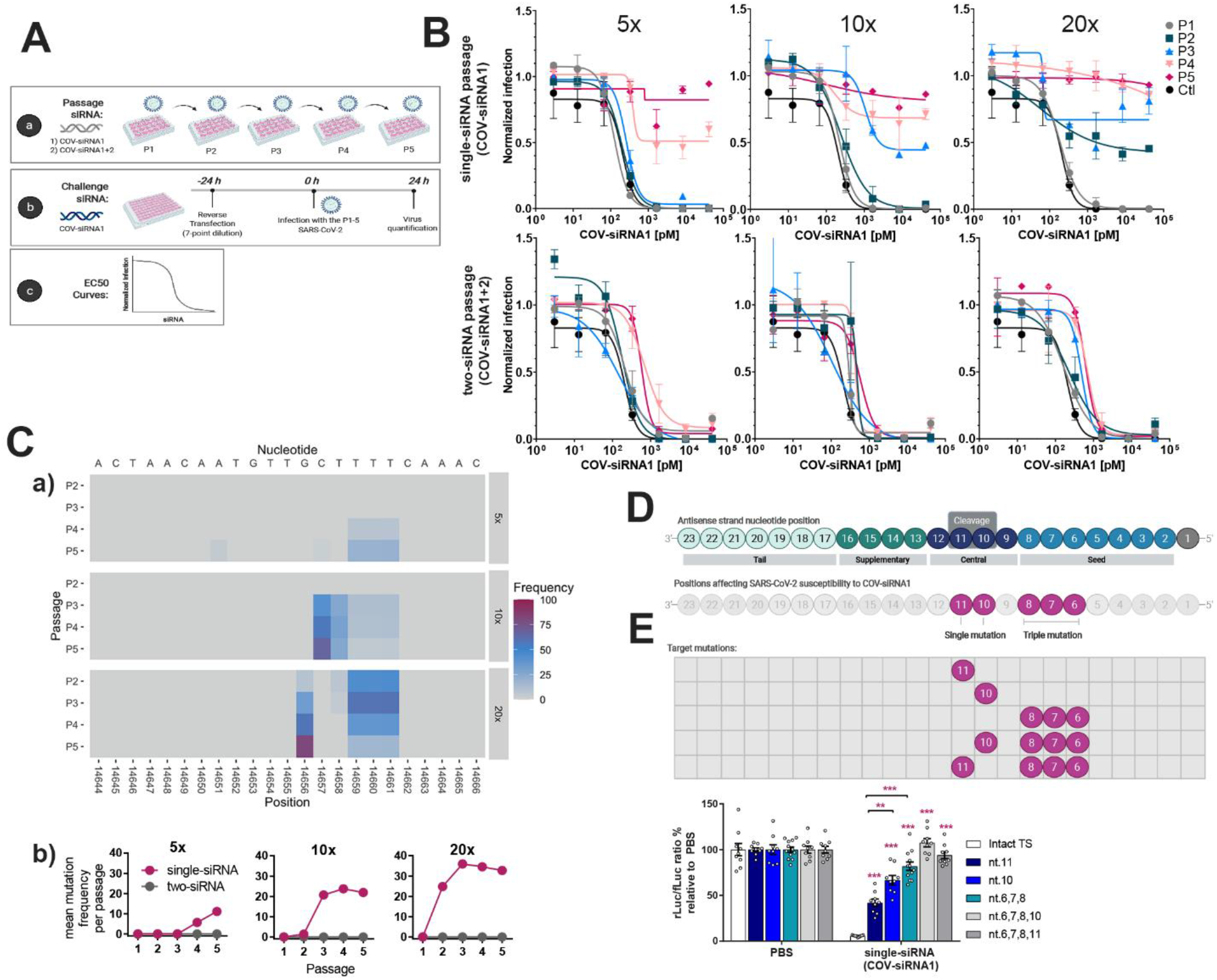
In Vitro Analysis of Viral Resistance to COV-siRNA1+2 in a SARS-CoV-2 Infection Assay. **(A)** SARS-CoV-2 was serially passaged in Vero E6 cells pre-treated with single COV-siRNA1 or COV-siRNA1+2 combination at 5x, 10x, and 20x EC_50_. (b) The virus-containing supernatants from each passage were then challenged with COV-siRNA1. **(B)** viral susceptibility to COV-siRNA1 was evaluated by EC_50_ determinations from single experiment with two independent repeats ± SD. **(C)** deep-sequenced at the siRNA target sites. Mutations identified with >2% frequency in samples treated with single-COV-siRNA1 are represented. No mutations were identified in samples treated with the COV-siRNA1+2 combination. **(D)** Schematic showing regions in the siRNA known to play different roles in RNAi activity (*top*) in relation to antisense strand positions impacted by observed viral resistance mutations resulting from COV-siRNA1 treatment (*bottom*). **(E)** *(top)* Dual-luc reporters containing either the intact viral target site (TS) of COV-siRNA1, or the target site mutated at a single or multiple nucleotide positions, as identified in the diagram. *(bottom)* Columns represent mean ± SEM luciferase (rLuc/fLuc) ratios following co-transfection of the indicated dose of COV-siRNA1 normalized to PBS-treated control. Each dot represents an individual replicate from three individual experiment with three repeats each. \*\**P*< 0.01, \*\*\**P*< 0.001 as determined by one-way ANOVA, Tukey post-hoc analysis.

### Identity of target site mutations after single siRNA treatment

To understand the reduced susceptibility observed following sequential treatment with a single siRNA, we deep-sequenced the SARS-CoV-2 genome at the siRNA binding site for all passages at all concentrations using a 2% frequency cutoff. Serial passaging with COV-siRNA1 alone led to the emergence of point-mutations in the COV-siRNA1 target site, but not in the COV-siRNA2 binding site **(Fig. 2C)**. The mutations at higher passages exist within the initial site of target RNA recognition and binding (the siRNA seed region). These mutations always appeared as a triple thymine mutation (TTT) at antisense nucleotides 6-8 (nt.6-8). At the highest passages, the triple mutation decreased, and two different mutations dominated at the predicted cleavage site of COV-siRNA1 (nt.10, nt.11). The extent and frequency of mutations correlates with the observed shifts in viral EC_50_. Both synonymous and non-synonymous base substitutions were seen at mutated sites with different doses of single siRNA treatment. In contrast, no mutations were detected at either siRNA target site at any viral passages when using the two-siRNA combination.

Mutations within the COV-siRNA1 target binding site, in both seed and central regions across the cleavage site, would be expected to compromise RNAi-mediated silencing activity (*11*). To investigate the underlying mechanism driving viral resistance, we evaluated COV-siRNA1 in a dual-luciferase reporter containing the intact viral target site (TS), as compared to those harboring single mutations within the cleavage region (nt.10, nt.11), the triple mutation at the seed region (nt.6-8), or a combination of mutations in both regions simultaneously **(Fig. 2C–D)**. As expected, treatment with COV-siRNA1 led to robust repression of reporter activity in the presence of the wild-type viral TS. In contrast, significantly reduced repression was observed when single mutations at nt.10 and 11 were introduced, consistent with the known positional base pairing requirements for RNAi-mediated target cleavage (*11*). We also observed a strong impact of the triple mutation in the seed region on COV-siRNA1 activity, highlighting the importance of complementarity in this region for RISC engagement and activity. We found that individual mutations at nt.6-8 positions had no significant impact on siRNA activity while double mutations were less tolerated, suggesting the need for more than one mutation within the seed region to significantly diminish siRNA efficacy **(Fig. S4)**. Overall, these results confirm that mutations observed with the single siRNA approach introduced vRNA-siRNA mismatches at key positions for RNAi-mediated activity. This highlights the need to employ a multi-siRNA combination approach to target distinct sites within the viral genome for therapeutic applications.

We performed an in-silico analysis evaluating the COV-siRNA1+2 target sequences across 4,386 SARS-CoV-2 sequences available in public databases to determine if any real-world mutations have appeared at these locations – including high interest variants Alpha, Beta, Gamma, Delta, Kappa, and Omicron **(Fig. 3)**. No mutations were observed within target sites of COV-siRNA1+2, indicating that these sequences have remained highly conserved across SARS-CoV-2 variants. Of note, many of the observed mutations in these variants of interest (VOI) and variants of concern (VOC) are in the spike gene within the receptor binding motif (RBM), a target of many antibody therapeutics and vaccine-induced immunity, resulting in ongoing concern around the potential loss of antiviral activity. With its distinct location of target sites, COV-siRNA1+2 may provide an orthogonal antiviral approach that remains agnostic to any newly arising mutations in the RBM.

**Fig. 3.**
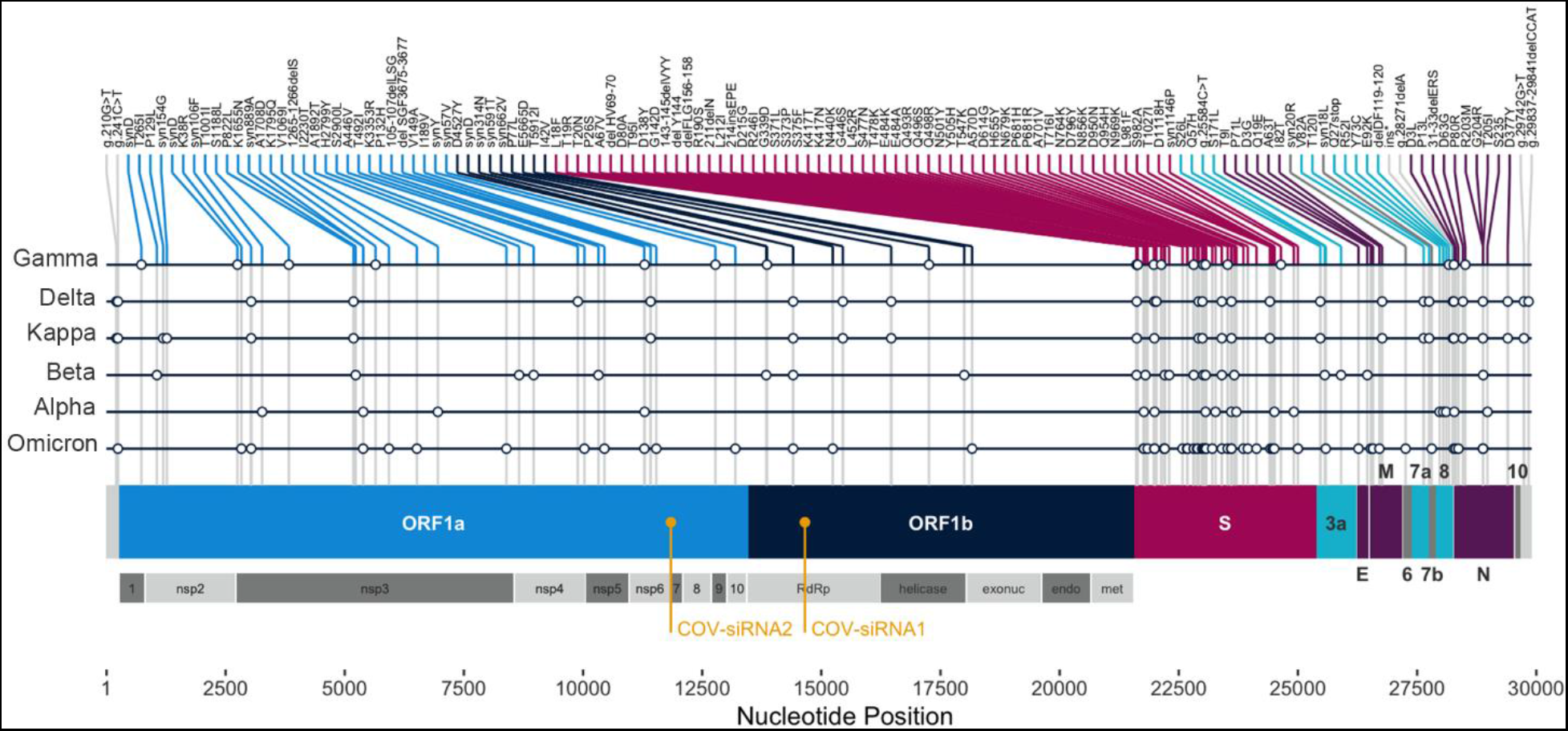
COV-siRNA1+2 target sites are conserved across SARS-COV-2 variants. Schematic derived from the in-silico analysis of COV-siRNA1+2 target sites relative to the mutations identified in the SARS-CoV-2 variants. Variations with a frequency greater than 50% were designated characteristic of the lineage. The coordinates and the specific changes caused by the mutation are labeled relative to the reference genome.

### COV-siRNA1+2 protects Syrian hamsters in a phenotypic model of SARS-CoV-2 infection

The in vivo efficacy of the siRNA combination was evaluated in the Syrian hamster model of SARS-CoV-2 infection (*12–14*). COV-siRNA1+2 (1, 10, 30 mg/kg) was administered IN 7 days before infection while siRNA targeting luciferase (30 mg/kg) was used as a non-specific control siRNA (**Fig. 4A and S5**). Two additional groups were treated IN one day after infection with 30 mg/kg of control siRNA or COV-siRNA1+2 (**Fig. 4B and S5**). At Day 0, animals were infected IN with SARS-CoV-2 (WA1). When administered prophylactically, COV-siRNA1+2 protected animals from body weight loss in a dose-dependent manner. At 7 days post-infection, hamsters pre-treated at day −7 with 1, 10, or 30 mg/kg IN of COV-siRNA1+2 showed 40%, 87%, and 104% of body weight recovery, respectively, relative to animals treated with control-siRNA **(Fig. 4A)**. In contrast, hamsters receiving 30 mg/kg of COV-siRNA1+2 one day after infection lost body weight at the same rate as control-siRNA treated groups **(Fig. 4B),** likely due to a delayed onset of activity consistent with the RNAi mechanism of action.

**Fig. 4.**
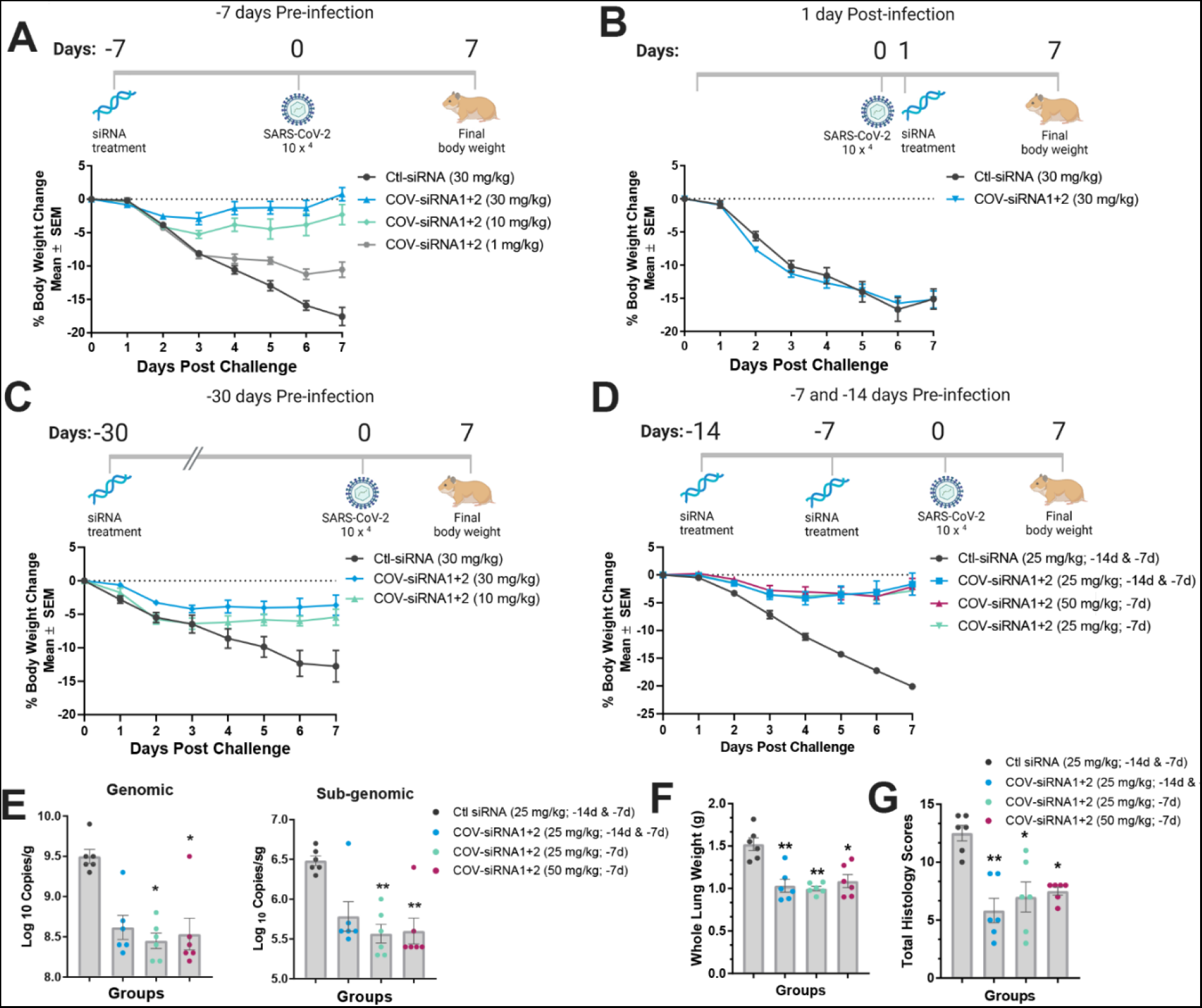
COV-siRNA1+2 efficacy in a hamster SARS-COV-2 infection model. Body weight change in hamsters treated with COV-siRNA1+2 combination or control siRNA **(A)** −7 days pre-infection, **(B)** +1-day post-infection, **(C)** −30 days pre-infection, and **(D)** −14 and/or −7 days pre-infection at the indicated concentrations (mg/kg) via intranasal instillation (IN). **(E-G)** Viral titers and histological examinations in lung tissues at 7 days post-infection of animals pre-treated with the siRNA combination at −14 and/or −7 days. **(E)** Each dot represents log _10_ genomic or subgenomic copy numbers of SARS-CoV-2 per individual animal by RT-qPCR. **(F)** Columns represent the mean whole lung weight per group or **(G)** mean lung histopathology score per group, graded from 1-5 depending on severity for, bronchioalveolar hyperplasia, alveolar hemorrhage, mononuclear cell vascular/perivascular inflammation, syncytial cells, mixed cell inflammation (bronchoalveolar, alveolar and/or interstitial), alveolar or perivascular edema, mesothelial hypertrophy and/or pleural fibrosis. Graphs show the means ± SEM; N=5-6; \**P*< 0.05, \*\**P*< 0.01 as determined by one-way ANOVA (Kruskal-Wallis test) followed by Dunn’s multiple comparison test.

The duration of the prophylactic benefit was also evaluated in animals treated with COV-siRNA1+2 at 30 days pre-infection. Results showed that hamsters were significantly protected from body weight loss, nearly to the same extent as was seen with a shorter pre-treatment interval **(Fig. 4A and Fig. 4C).** Durability is consistent with that observed with other RNAi therapeutics in preclinical species (*15*). We also compared a single IN dose of 25 mg/kg or 50 mg/kg at day −7 vs. two sequential IN doses of 25 mg/kg at day −14 & −7 **(Fig. 4D)**. These data suggest that 25 mg/kg provides the highest achievable protection with COV-siRNA1+2 in this model.

Viral burden and lung pathology were evaluated in animals treated prophylactically at day −7 and/or −14. Lung tissues were evaluated for genomic and sub-genomic RNA **(Fig. 4E)**. These data show significant reductions of viral genomic and sub-genomic RNA in groups treated with COV-siRNA1+2. Lung weights of treated animals were also significantly lower than control groups, suggesting reduced lung inflammation **(Fig. 4F)**. Histological evaluation of lung tissue was performed with pathology findings graded from 1-5 depending on severity. Total scores of microscopic findings were calculated **(Fig. 4G; Table S3)** and representative images were obtained **(Fig. S6).** Animals treated with all doses of COV-siRNA1+2 exhibited decreased severity or incidence of SARS-CoV-2-related microscopic findings of bronchioloalveolar hyperplasia, hemorrhage, inflammation, syncytial cell, and edema, except for pleural fibrosis when compared to control animals.

Taken together, these data show that prophylactic COV-siRNA1+2 treatment provides durable and robust protection in a preclinical model of SARS-CoV-2 infection.

## Discussion

We identified multiple siRNAs spanning the SARS-CoV-2 genome with potent antiviral activity when tested in vitro using a SARS-CoV-2 infection model, alone or in combination. One of the key benefits expected with the combination RNAi approach is minimizing viral resistance as it would require the simultaneous emergence of mutations in two distinct regions of the viral genome. COV-siRNA1+2 targets highly conserved sites on the ORF1ab. The conservation suggests that those regions are likely critical for viral fitness and less likely to mutate. The COV-siRNA1 binding site lies within the NSP12 region, which encodes the RNA-dependent RNA polymerase (RdRp), critical for viral replication and a target of small molecule antivirals such as molnupiravir and remdesivir. The target site does not overlap with regions encoding the RdRp active sites or known remdesivir or molnupiravir resistance mutations, thus no treatment interference is expected (*16–18*). The target site of COV-siRNA2 is in the junction between NSP 6/7, encoding additional cofactors of the replicase.

We first evaluated COV-siRNA1 as a single siRNA in an in vitro serial-passaging study to examine the potential for viral resistance to develop. After five passages, we observed the emergence of resistant virus populations. Resistance occurred sooner at higher concentrations, likely due to greater selective pressure. To address this limitation, we added a second siRNA (COV-siRNA2) as a two-siRNA combination and found this could prevent the emergence of escape mutants. This is consistent with prior results demonstrating that the use of multiple siRNAs to target other CoVs was more effective than a single siRNA (*19*). Using deep sequencing and a luciferase reporter assay, we confirmed that viral mutations observed with the single siRNA approach accumulated vRNA mismatches at key positions causing a reduction of siRNA efficacy. This finding highlighted the need to employ a multi-siRNA targeting distinct sites or combinations of orthogonal interventions to minimize the risk of viral resistance in any real-world therapeutic applications.

Since late 2020, new SARS-CoV-2 variants have steadily replaced the spike D614G variant highly prevalent in the first wave of the pandemic. We performed an exhaustive in silico evaluation of current WHO VOI and VOC including those encoding the receptor binding motif N501Y spike mutation which emerged simultaneously in the variants Alpha, Beta, Gamma, Delta, and Omicron (*20–22*). While these variants encode substitutions across the viral genome, they possess similar mutation patterns associated with improved affinity for the ACE2, the known entry receptor for SARS-CoV-2. Favorably, the sequences targeted by COV-siRNA1+2 are located far away from the spike encoding region. Our in-silico analysis of public databases suggests no anticipated loss of activity (target site sequence fidelity) with any known VOC or VOIs.

The Syrian hamster is a well-established animal model of SARS-CoV-2 infection and has been used in the development of vaccines and antibodies (*12*, *13*). It supports infection due to its high ACE2 receptor homology to human; infected animals manifest clinical symptoms including body weight loss and histopathological changes in the lung. We demonstrated that prophylactic administration of COV-siRNA1+2 provided robust protection to hamsters, consistent with a significant reduction of viral loads and reduction in the severity of clinical signs. We confirmed the durability of protection with dosing 1-month prior to infection challenge. The extent of activity was consistent with previous studies using other siRNAs, demonstrating 56 days of sustained silencing of an endogenous lung target in rodents, employing the C16-conjugation chemistry (*8*). This suggests a long duration of activity for C16-siRNAs in the lung, similar to what we have seen previously with GalNAc-siRNAs in the liver (*23*).

We also evaluated COV-siRNA1+2 in a post-infection treatment paradigm where animals were dosed one day after viral challenge, and we did not observe therapeutic benefit. This is consistent with a delayed onset of action for RNAi along with rapid onset and resolution of infection in the hamster model.

There are multiple settings where prophylactic treatment with COV-siRNA1+2 could have utility. One could consider use in combination with vaccines to provide a stronger antiviral barrier in high-risk populations, such as the elderly, those with comorbidities, front-line workers, or in immunocompromised status where vaccines are less effective. Due to the high conservation of the target sites, COV-siRNA1+2 may retain activity against yet unknown coronaviruses that may emerge in the future and could be rapidly developed to combat future pandemics.

COV-siRNA1+2 has been designed to be suitable for *in vivo* investigation. The siRNA modifications within COV-siRNA1+2 have been part of multiple approved RNAi therapeutics and the C16 conjugate has been studied extensively in preclinical species and is being used in a development candidate targeting amyloid precursor protein entering investigational clinical trials in 2022 (*8*).

Taken together, these data indicate that COV-siRNA1+2 could be a key tool to prevent COVID-19, with a unique profile compared to vaccines, monoclonal antibodies, and other therapies. COV-siRNA1+2 could serve as a potentially long-acting prophylactic agent to protect from severe disease and join the revolution of RNA-based therapeutics fighting current and future pandemics.

## Acknowledgments

The authors thank the High-throughput Synthesis Team; Ivan Zlatev, Klaus Charisse, and the Medium Scale Synthesis Team; Ligang Zhang and the CMC Team for siRNA synthesis; Alex Eaton and MaryBeth Kim from the Research Lead Discovery team, and Jeffrey Zuber from the Bioinformatics team. We would like to acknowledge Swagata Kar and the Bioqual team for their in vivo support. We thank Kevin Sloan and Kevin Dooley for reviewing the manuscript. Illustrations were created with BioRender.com.

## Author contributions

Conceived studies: YIAR, FAL, JM, MKS, CRB, DJF, JD, SC, JN, AS, MAM, GH, CMH, AA, VJ. Designed studies and experiments: YIAR, FAL, JM, MKS, CRB, DJF, ABC, TN, MS, AS, RS, SSM, JDB, DB, DW, MM, AR, JD, SC, JN, AS, KS, SH, HWV, KF, MAM, GH, CMH, AA, VJ. Performed experiments: YIAR, FAL, JM, TN, MS, MMR, HK, AS, RS, SSM, SL, AK, TR, MM. Analyzed and interpreted data: YIAR, FAL, JM, MKS, CRB, DJF, TN, MS, MMR, AS, AS, RS, JDB, DB, AK, DW, MM, AR, JD, SC, AS, KS, SH, HWV, KF, MAM, GH, CMH, AA, VJ. Supervised: KF, MAM, GH, CMH, AA, VJ. Prepared the manuscript with input from all authors: YIAR, VJ

## Competing interests

All authors were employees of Alnylam Pharmaceuticals or Vir Biotechnology with salary and stock or stock options when the work was conducted. HWV is a founder of PierianDx and Casma Therapeutics. Neither company provided funding for this work or is performing related work.

## Data and materials availability

All data are available in the main text or the supplementary materials.

## Supplementary Materials

Materials and Methods

Figs. S1 to S6

Table S1 to S3

## Materials and Methods

### Sequence Analysis for selection siRNA binding sites

Paired-end reads were trimmed to 2X150 upon inspection with FastQC to retain segments of high quality (>Q30). Illumina adapters were clipped using Trimmomatic. Read alignment to SARS-CoV-2 Wuhan-Hu-1 reference (NCBI: NC_045512.2) was performed with Burrows-Wheeler Aligner (BWA). Variants were called with LoFreq* upon indel realignment and base quality recalibration, using a frequency threshold of 1%. Primers were excluded from variant calling. A consensus sequence was generated, mapped back to reference sequence coordinates and used to align reads a second time. The incorporation of variants from the first mapping iteration into a consensus sequence facilitates read alignment in case of mismatches with the reference. Variants were called again against the reference sequence from reads mapped to the consensus sequence. A third iteration of consensus generation read alignment and variant calling was performed. The variant call set from this last iteration was retained as final. Nucleotide variants were annotated with SnpEff to generate amino acid variants. Extensive QCs were performed at read, alignment and variant level using FastQC, samtools, picard, mosdepth, bcftools and MultiQC. An end-to-end workflow was automated using NextFlow. All programs are available through the Bioconda Initiative (bioconda.github.io).

The siRNA sequences directly targeting the SARS-CoV-2 genomic RNA were generated from the exemplar sequence in NCBI Genbank®. A custom script was used to identify stretches of the exemplar genome also present as complete matches in the full-length sequences of all available MERS-CoV (16), SARS-CoV (14), and SARS-CoV-2 (57) genome sequences from NCBI and EpiCov as of January 2020, in an effort to increase the likelihood of retaining full cross-reactivity as SARS-CoV-2 evolves. The script identified 1511 conserved SARS-CoV-2/SARS-CoV-1 19-nucleotide sequences as well as 7 SARS-CoV-2/MERS-CoV sequences. A subset of 349 unique sequences was selected for further evaluation.

### Oligonucleotide synthesis

All oligonucleotides were synthesized on an MerMade 192 synthesizer according to previously published protocols (*24*). 2’-OMe and 2’-F phosphoramidites were purchased commercially and used as previously reported. 2’-*O*-C16 phosphoramidites (*25*) were used at a concentration of 100 mM in acetonitrile with no other changes to synthetic protocols.

### Bioinformatic Design of Reporter Plasmid Design for siRNA Screening

The synthesized siRNA duplexes were screened for target reduction relative to mock transfection in a psiCHECK2 dual-luciferase reporter assay that incorporated a concatemer insert designed such that all targets of the candidate siRNA duplexes were represented (**Fig. S1C**). The construct also contained a 23-mer target sequence that served as a positive control target site with a previously well-characterized siRNA. Randomized concatemers were analyzed for RNA secondary structure using the RNA structure software toolkit (*26*). Two concatemers for (+) RNA genome and one for (-) RNA were selected based on the predicted accessibility of the siRNA target sites at each position and were complementary such that slightly lower accessibility for a given target in one plasmid was compensated for by improved accessibility in the alternate plasmid. The SARS-CoV-2 concatemeric sequences were synthesized and incorporated individually into the psiCHECK2 vector by Blue Heron Biotech [Bothell, WA]. The final vectors are referred to as psiCHECK2-CoV-2A, psiCHECK2-CoV-2B, and psiCHECK-negCoV2.

### Dual-luciferase assays

Cos7 cells were co-transfected with any of the reporter plasmids, and siRNAs in 384-well plates format at a density of 5×10^4^ cells per well. Forty-eight hours after transfection, Firefly (transfection control) and *Renilla* (fused to the SARS-CoV-2 target sequence) luciferase were assayed using the Dual-Glo™ Luciferase Assay System. The siRNAs with ≥80% target reduction relative to mock-transfected controls in both psiCHECK-CoV2 plasmids were selected for further evaluations. For mutational analyses, a series of psiCHECK2 reporter plasmids were constructed, in which a single 3’UTR copy of either the intact or mutated 23-nucleotide viral target site of COV-siRNA1 was cloned downstream of *Renilla* Luciferase. The siRNA activity was determined by normalizing the *Renilla* (target) signal to the firefly (control) signal within each well. The magnitude of siRNA activity was then assessed relative to cells that were transfected with the same vector but were not treated with siRNA. All transfections were performed at least in triplicate.

### SARS-CoV-2 in vitro infection assay

Vero E6 cells were reverse transfected with the corresponding concentration of siRNA in a 96-well plate format using RNAiMax. Control wells were either mock transfected (no siRNA) or transfected with an siRNA targeting the luciferase ORF (control siRNA). Cells were infected with SARS-CoV-2 (USA-WA1/2020, passage 2) (BEI; NR-52281) at MOI 0.001 or MOI 0.01 at 24 h post transfection. Four hours post infection, inoculum was removed, and cells were rinsed with phosphate-buffered saline (PBS). Supernatant was removed at 24 h (MOI 0.01) or 48 h (MOI 0.001) post infection and RNA used was used for RT-qPCR evaluation and while fixed cells were used for IFA quantification.

### Quantification of extracellular SARS-CoV-2 genomes by RT-qPCR

Viral RNA was extracted from the cell culture supernatant using the NucleoSpin 96 Virus kit (Macherey-Nagel). Quantification of viral genomes was performed using the Luna Universal Probe One-Step RT-qPCR Kit (New England Biolabs) with a primer/probeset binding in the orf1ab region (forward: CCCTGTGGGTTTTACACTTAA, reverse: ACGATTGTGCATCAGCTGA, probe: CCGTCTGCGGTATGTGGAAAGGTTATGG). A standard curve of defined dilutions of a synthetic SARS-CoV-2 RNA (Twist Bioscience) was used for normalization.

### Immunofluorescence analysis (IFA)

Cells were fixed with 4 % paraformaldehyde for 30 min, followed by two PBS washes and permeabilization with 0.125% Triton X-100 in PBS for 30 min. After blocking in 2% milk powder/PBS for 30 min, cells were incubated with a primary antibody targeting SARS-CoV-2 nucleocapsid protein (Sino Biological) at a 1:3000 dilution for 1h followerd by incubation with a secondary AlexaFluor647-labeled antibody. Nuclei were stained using Hoechst33342. Single images were acquired using an Echo Revolve inverted fluorescence microscope. IFA staining in whole wells was quantified using automated image acquisition, on a Cytation 5 Cell Imaging reader with 12 images per well to cover the complete well. Nuclei and AlexaFluor647-positive cells were counted using the manufacturer’s provided software. After subtraction of background (uninfected) controls, all signals were normalized to the mock transfected controls.

### Resistance selection using fixed concentrations

Resistance selection using fixed concentrations of COV-siRNA1 or COV-siRNA1+2 was conducted in Vero E6 cells cultured in DMEM supplemented with 10% FBS (VWR) and 1x Penicillin/Streptomycin (Thermo Fisher Scientific). 1.2 x 10^5^ Vero E6 cells per well in a 24-well plate were reverse transfected with 5x, 10x or 20x EC_50_ of COV-siRNA1 (330 pM, 660 pM or 1320 pM, respectively) or of an equimolar combination of COV-siRNA1 and COV-siRNA2 at the same combined total concentrations. As a non-targeting control, cells were reverse transfected with 1320 pM of a control siRNA targeting the luciferase gene. All transfections used RNAiMax (Thermo Fisher Scientific) according to the manufacturer’s instructions. Six hours post reverse transfection, cells were incubated with SARS-CoV-2 (USA-WA1/2020, passage 2) at an MOI of 0.01 in infection medium (DMEM, 10% FBS) for 1 h at 37°C. After viral adsorption, cells were washed with DMEM and overlaid with infection media. For each condition, three independent wells were transfected and infected, and the supernatants were pooled together at the time of harvest. Infected cells were monitored visually for cytopathic effect (CPE) daily. When the cells in the luciferase-targeting control wells exhibited > 50% CPE, the culture supernatants were harvested, diluted 1:10 and added to fresh Vero E6 cells in 24-well plates that had been reverse-transfected with the equivalent amounts of siRNA(s) as used for the initial passage. Selection continued for a total of 5 passages. At each passage, supernatant was aliquoted and frozen at −80°C for further analyses.

### Determination of Viral Titer by Focus-Forming Assay

Viral titers were determined using a focus-forming assay (FFA) on Vero E6 cells. One day prior to infection, 1.5 x 10^4^ Vero E6 cells were plated in black-walled, clear bottomed 96-well plates. The next day, undiluted or 4-point 10-fold serially diluted virus samples using 10% FBS-containing media were adsorbed onto cells for one hour at 37°C. The cells were washed once and overlaid with 1% methylcellulose (Sigma-Aldrich) in serum-containing medium. At 24 h post-infection, the methylcellulose overlay was removed, and cells were washed with PBS. Cells were fixed with 4% paraformaldehyde (PFA), incubated for 30 minutes at RT, then washed with PBS. The cells were permeabilized with 100 μL of 0.25% Triton X-100 (Sigma-Aldrich) in PBS for 30 minutes at RT. The Triton X-100 was removed, cells were washed twice with PBS, and incubated with 50 μL of SARS-CoV-2 nucleocapsid (N) antibody (Sino Biological) at 1:2,000 in blocking buffer (2% milk powder/PBS) for 1 hour at RT. Plates were washed three times with PBS and then incubated for 1 hour at RT with 50 μL/well of goat anti-rabbit-Alexa647 (Thermo Fisher Scientific) secondary antibody at 1:1,000 in blocking buffer along with 1 μg/ml Hoechst 33342 (Thermo Fisher Scientific). After washing three times with PBS, plates were imaged on a Cytation5 (BioTek) plate reader (12 images at 4X magnification) with fluorescence detected in DAPI (377,447nm) and Cy5 (628,685nm) channels. Nucleocapsid-positive foci were counted from images and used to determine focus-forming units/mL supernatant (FFU/mL).

### Evaluation of Antiviral Activity Against Selected Virus

Cell supernatants from viral passages containing detectable virus as determined by FFA were evaluated for a shift in EC_50_ values in an siRNA antiviral activity assay. 7-point 5-fold serial siRNA dilutions prepared in PBS and Vero E6 cells were reverse transfected in 96-well plates with each dilution in duplicates (range: 41250 to 3 pM final concentration). Twenty-four hours later, cells were infected with 20 FFUs of the virus-containing cell supernatants generated during passaging. The viral inoculum was removed after 1 h and media was replaced. At 24 h post-infection, the cells were fixed with 4 % PFA and viral nucleocapsid protein was immunostained as described above.

### Sequencing of siRNA binding sites within the SARS-CoV-2 genome

#### RNA isolation

300 μL of cell supernatant was added to 900 μL of Trizol and stored at −80°C for further analysis. Trizol collected samples were subjected to RNA isolation using PureLink RNA Mini Kit with the incorporation of on-column PureLink DNase Treatment. Reverse transcription reactions were performed with 6 μL of purified RNA and random hexamer primers using the NEB ProtoScript II First Strand cDNA Synthesis kit. The resulting cDNA was used as a template for PCR amplification of the siRNA binding sites, two primer pairs per site, using KapaBiosystems polymerase (KAPA HiFi HotStart ReadyMix) with primers, 5’-acgctgcttctggtaatc-3’ plus 5’-agaaacccttagacacagc-3’, and 5’-ctagataaacgcactacgtg-3’ and 5’-cctgagcaaagaagaagtg-3’ for the COV-siRNA1 binding site, and 5’-agatgccttcaaactcaac-3’ plus 5’-ttctactctgagttgttgc-3’, and 5’-ttggtggcaaaccttgtatc-3’ plus 5’-tgtgtaactggacacattg-3’ for the COV-siRNA2 binding site. Each PCR with a primer pair was carried out independently, then PCR products from each treatment were pooled for purification and subsequent library preparation. Amplification conditions included an initial 3 min at 95°C, followed by 24 cycles with 20 sec at 98°C, 15 sec at 60°C and 72°C for 15 sec, with a final 1 min at 72°C. Pooled PCR products were purified using AMPure XP beads. The size of the amplicon was confirmed by analyzing 2 μL of PCR products using the Agilent D1000 ScreenTape System. Products were quantified by analyzing 1 μL with the Quant-iT dsDNA High-Sensitivity Assay Kit. Fifty ng of purified PCR product was used as input for library construction using the NEBNext Ultra II DNA Library Prep kit. NEBNext Multiplex Oligos for Illumina Dual Index Primer Set 1 was used for library construction, with a total of 4 PCR cycles. Libraries size was determined using the Agilent D1000 ScreenTape System and quantified with the Quant-iT dsDNA High-Sensitivity Assay Kit. Equal amounts of each library were pooled together for multiplexing and ‘Protocol A: Standard Normalization Method’ of the Illumina library preparation guide was used to prepare 8 pM final multiplexed libraries with 1% PhiX spike-in for sequencing. The MiSeq Reagent Kit v3 (600-cycle) was used for sequencing the libraries on the Illumina MiSeq platform, with 150 cycles for Read 1, 150 cycles for Read 2, 8 cycles for Index 1, and 8 cycles for Index 2.

### Variant Analysis Protocol for SARS-CoV-2 Iota and Omicron Variants

Visualizations for characteristic polymorphisms of SARS-CoV-2 lineages were produced based of annotations published in papers that provided a listing of those mutations. For the Iota and Omicron variants, no such detailed references were available, and the data was obtained through analysis of SARS-CoV-2 genomic sequence data downloaded from the EpiCoV data hosted by the GISAID Initiative. The FASTA-formatted sequence data and associated tab-delimited metadata table for the Iota variant were downloaded on 24 February 2021 from the epicov.org web site (*27*); this data represented a current snapshot of the repository’s SARS-CoV-2 sequence data as of that date. The FASTA sequence data was indexed using the makeblastdb program from the ncbi-blast software package (*28*). A custom R script was written to load the associated metadata table and subset the records that met the following criteria: PANGO lineage = “B.1.526”, Host = “Human”, length > 29500. The corresponding 593 lineage-specific sequence records were extracted from the larger sequence set using the blastdbcmd program to a separate file.

Multiple sequence alignment of the lineage-specific sequences was performed using MAFFT(*29*) to align the sequences as fragment alignments (*30*) the SARS-CoV-2 reference genome (NCBI accession NC_045512.2) with the parameters --6merpair --thread −1 --addfragments lineage-B.1.526.fasta NC_045512.2.fasta. The MAFFT application output is a multiple sequence alignment of the lineage sequences relative to the reference genome. The multiple sequence alignment file was processed using a custom R script and the BioConductor Biostrings library to parse the FASTA alignment file and produce a table of insertions, deletions, and substitutions in the lineage sequences relative to the reference by simple pair-wise comparison of each aligned lineage sequence to the reference. The frequency for each individual variation was computed using the Biostrings consensusMatrix() call to compute the frequency of the most prominent variation at each physical position. Given the positions relative to the genomic reference sequence, corresponding positions and peptide changes were computed based off the feature annotation for the NCBI sequence record for the reference genome. Variations with a frequency greater than 50% were designated characteristic of the lineage. The coordinates and the specific changes caused by the mutation were used to label a diagram of the mutations relative to a schematic of the reference genome and its genes.

For the Omicron variant, the same procedure was performed using EpiCoV database snapshot from 27 November 2021 and the criteria: PANGO lineage = “B.1.1.529”, Host = “Human”, length > 29500. These criteria identified 77 genomic sequence entries. The MAFFT parameters were --6merpair --thread −1 --addfragments lineage-B.1.1.529.fasta NC_045512.2.fasta.

### SARS-CoV-2 infection in Syrian Golden Hamsters

#### Ethics Statement and Animal Exposure

Animal research was conducted under BIOQUAL Institute Institutional Animal Care and Use Committee (IACUC)-approved protocols, 20-152 in compliance with the Animal Welfare Act and other federal statutes and regulations relating to animals and experiments involving animals. BIOQUAL is accredited by the Association for Assessment and Accreditation of Laboratory Animal Care International and adheres to principles stated in the Guide for the Care and Use of Laboratory Animals, National Research Council. Animals were monitored at least twice daily, and enrichment included commercial toys and food supplements. Prior to all blood collections, animals were anesthetized using Ketamine/Xylazine. At the end of the study, animals were euthanized with intraperitoneal overdose of sodium pentobarbital.

#### siRNA dosing and SARS-CoV-2 challenge

Syrian Golden hamsters approximately 6-8 weeks of age, male (N=5-6 per group) were dosed intranasal (IN) with a luciferase siRNA (Ctl-siRNA) or COV-siRNA1+2 at Day −7, Day-14, or Day −30 prior to SARS-CoV-2 infection. Two groups were given the siRNAs via IN dosing one day after infection. Appropriate dose was administered with a total volume of 100 μL per animal (50 μL/nostril). Hamsters were challenged IN on day 0 with SARS-CoV-2 administered dropwise with a pipet with a total volume of 100 μL per animal (50 μL/nostril) at a challenge dose of 3 × 10^4^ PFU per hamster. Animals were observed twice daily for clinical signs and body weights were measured on Day 0 prior to challenge and daily for 7 days. Terminal lung tissue were collected on day 7. Samples were analyzed by RT-qPCR targeting viral genomic and sub-genomic RNA using quantitative real-time PCR methods per Bioqual internal methodology.

#### Histology

At necropsy, left lung lobe was collected on day 7 and placed in 10% neutral buffered formalin for histopathologic analysis. Lung was processed to hematoxylin and eosin (H&E) stained slides and examined by a board-certified pathologist. Histopathologic findings were graded from 1-5, depending upon severity. Findings were scored as follows: Grade 0 – absent, Grade 1 – minimal (<10% of tissue affected); Grade 2 – mild (11–25% of tissue affected); 3 – moderate (26–50% of the tissue affected); 4 – marked (51–75% of the tissue affected); and 5 – severe (>75% of tissue section affected).

### Statistical Analysis

Statistical analyses were performed where indicated using Prism v7.01 (GraphPad Software). P values <0.05 were considered significant. Data were analyzed by one- or two-way analyses of variance (ANOVAs) followed by post hoc Tukey’s test or Dunnett’s test for multiple comparisons. Non-normally distributed data were log-transformed.

## Supplemental Figures

**Fig. S1.**
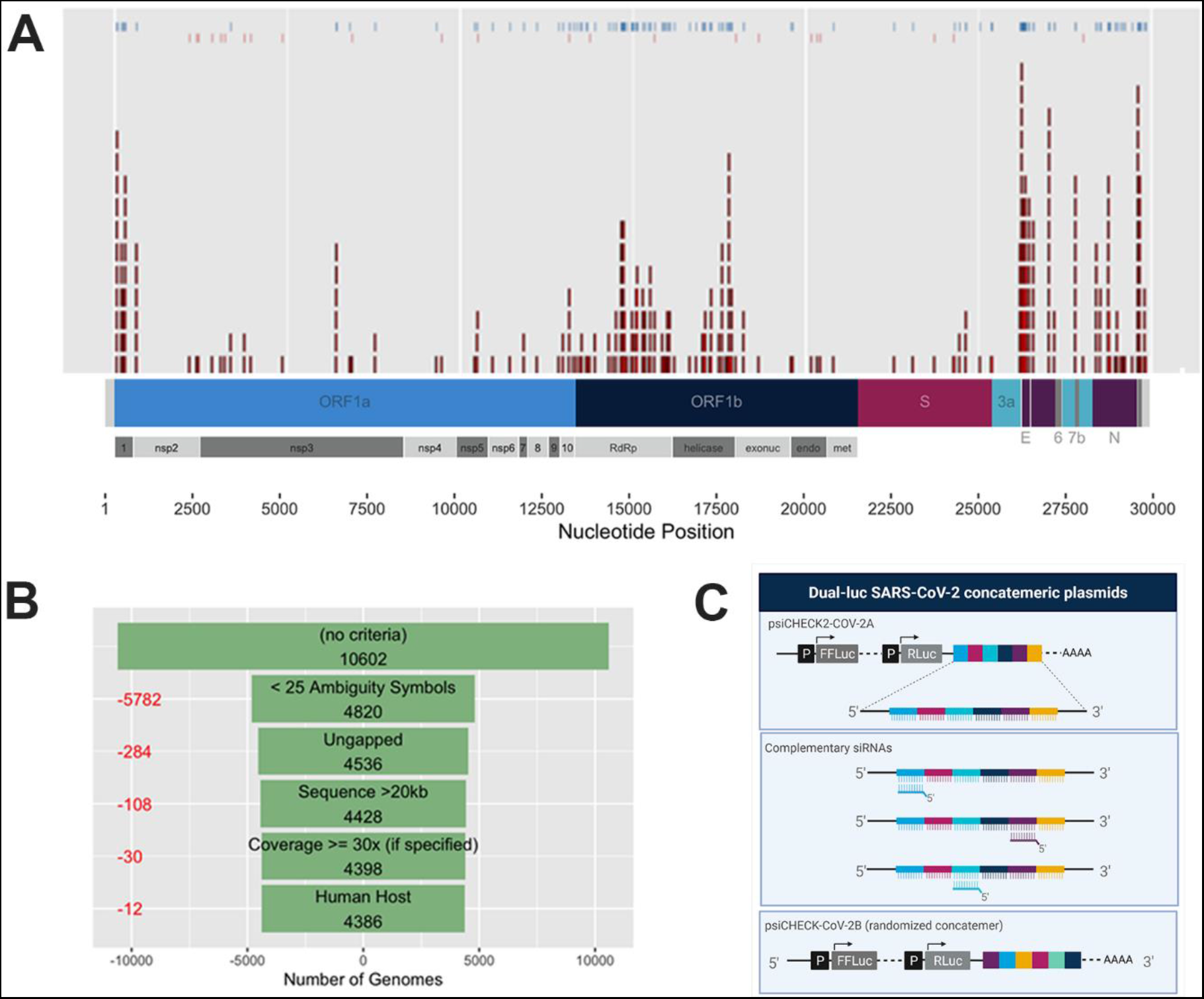
Bioinformatic evaluation of SARS-CoV-2 genome, target site selection and concatemeric plasmids illustration. **(A)** Genome binding sites of the selected siRNAs. Top ticks show siRNAs cross-reactivity to SARS-CoV (blue) or SARS-CoV + MERS (red). **(B)** Criteria on high-confidence genome selection from databases. **(C)** Dual-luciferase reporter concatemeric plasmids design.

**Fig. S2.**
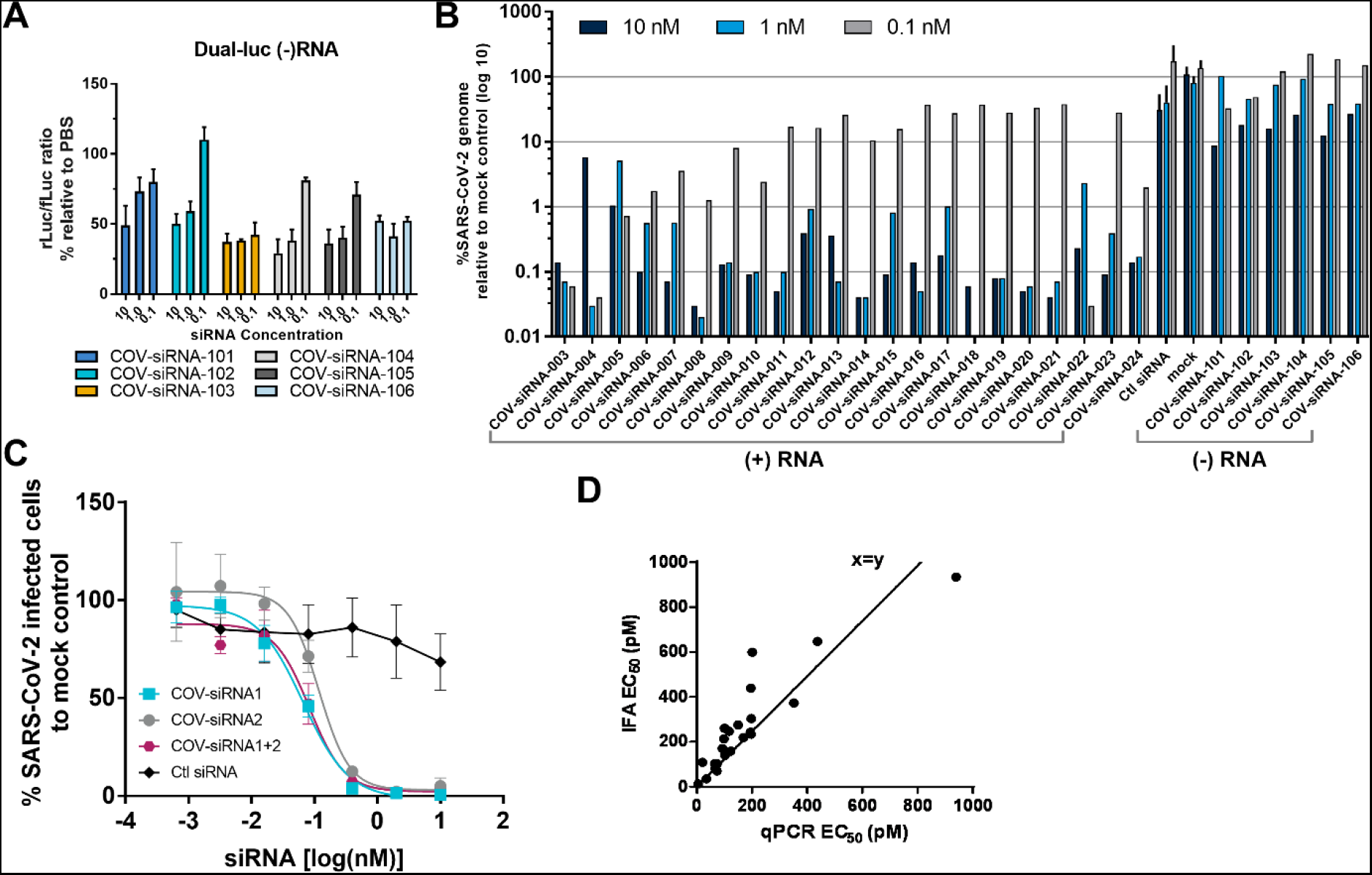
Additional in vitro screening data. **(A)** *Luciferase in vitro screening of (-) RNA SARS-CoV-2 targeted siRNAs*. Each column represents the mean normalized level of luciferase reporter activity (rLuc/fLuc) relative to PBS in psiCHECK-negCoV2 reporter plasmid from a single experiment with three independent repeats ± SD. Cos7 cells were reverse transfected at the indicated concentration of siRNA. **(B)** Vero E6 cells were reverse transfected with siRNA or controls and infected with SARS-CoV-2 isolate USA-WA1/2020. Twenty-four hours post transfection, cells infected, and supernatant was evaluated by RT-qPCR for viral genomic RNA from single experiment representative of one-four independent repeats ± SD **(C)** Dose-dependent knockdown of viral genomic RNA after siRNA treatment quantified by RT-qPCR from two independent experiment with three independent repeats ± SEM. **(D)** Correlation observed for both viral mRNA and N protein detection assay. Each dot represents an individual siRNA evaluated by both assays.

**Fig. S3.**
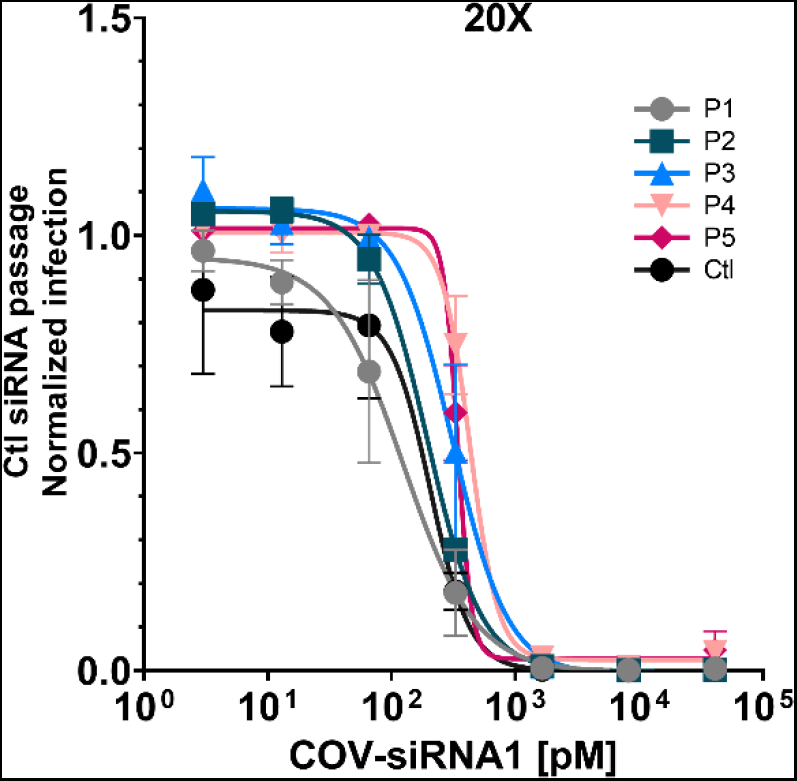
EC_50_ determinations to COV-siRNA1 after serial passage with control siRNA at 20x EC_50_. SARS-COV-2 virus obtained from all passages was susceptible to COV-siRNA1 and no EC_50_ curves shifts were observed. Graph shows EC_50_ determinations from single experiment with two independent repeats ± SD.

**Fig. S4.**
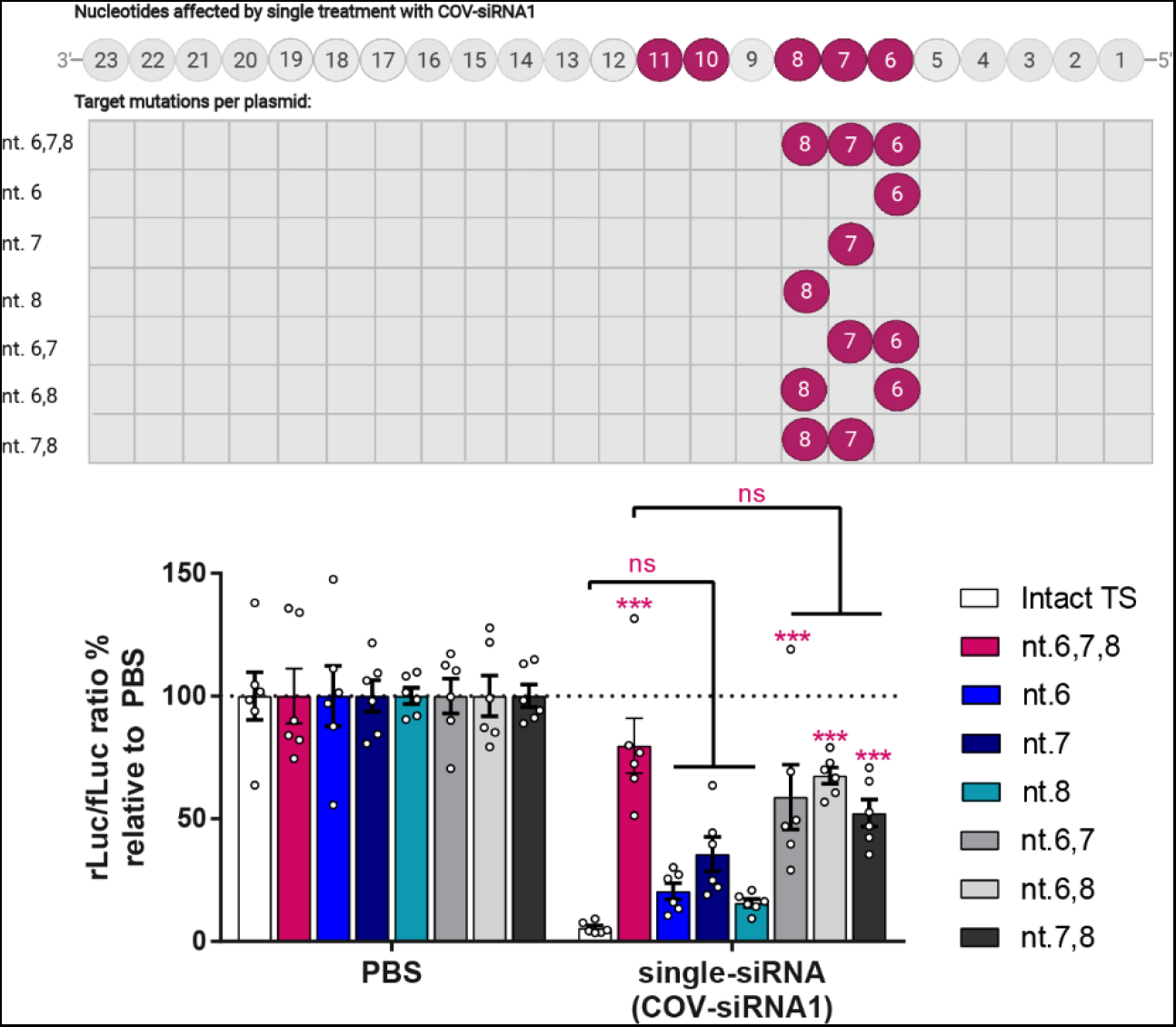
Target mutational assay relative to nucleotides 6-8 of the antisense strand of the COV-siRNA1. **(A)** Diagram showing target site composition of dual luciferase vectors containing single, double, or triple mutations in seed-matched region of viral target site (TS), relative to COV-siRNA1 antisense strand nucleotides (nt) 6-8. **(B)** Columns represent mean ± SEM luciferase (rLuc/fLuc) ratios following co-transfection of COV-siRNA1 normalized to PBS-treated control. Each dot represents an individual replicate from three individual experiment with three repeats each. ***P< 0.001, no significant (ns), as determined by one-way ANOVA, Tukey post-hoc analysis.

**Fig. S5.**
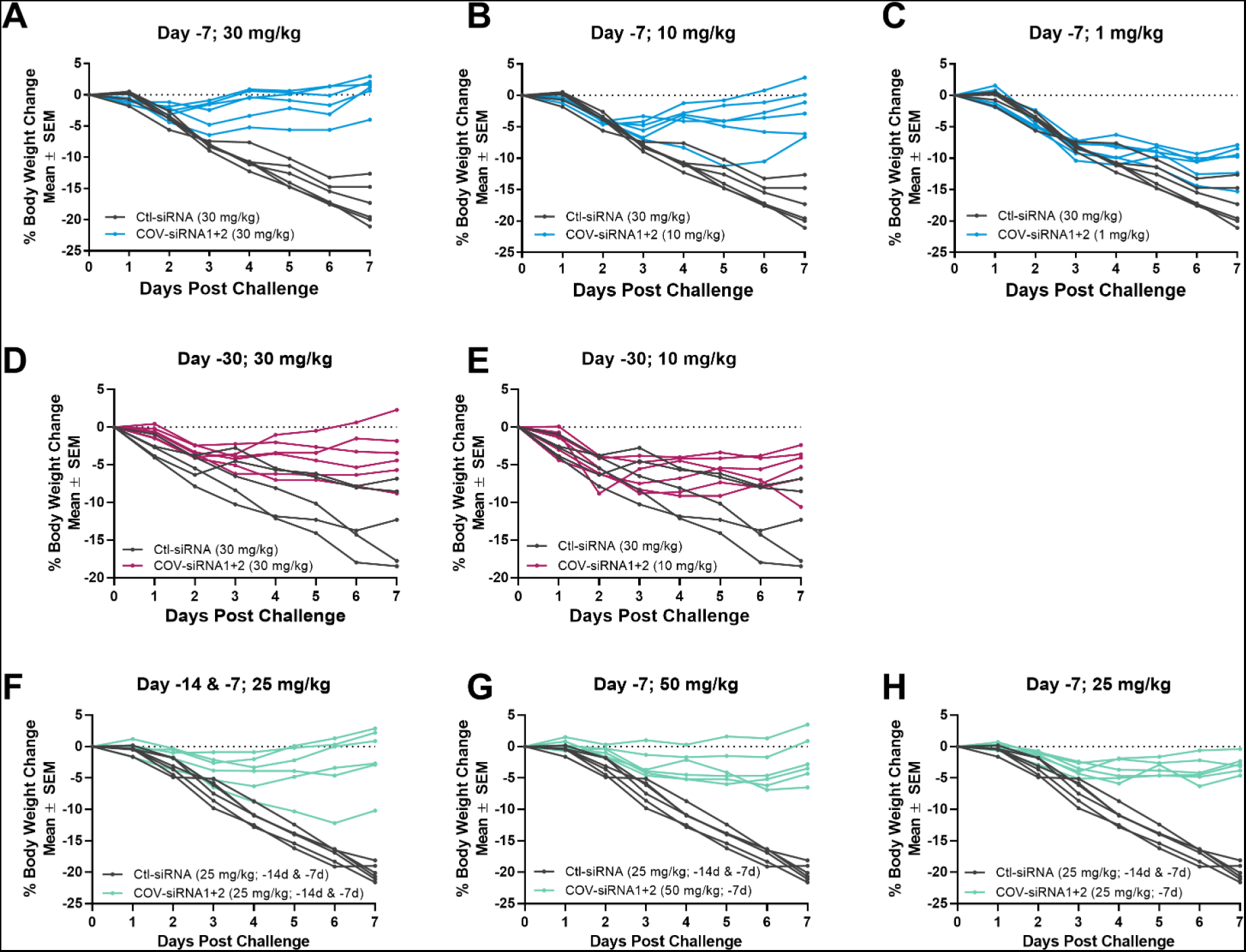
Body weight in single animals. Graphs show body weight change in hamsters treated with COV-siRNA1+2 combination or control siRNA at **(A-C)** −7 days pre-infection, **(D-E)** −30 days pre-infection, **(F-H)** −14 and/or −7 days pre-infection. At the indicated concentrations (mg/kg) via intranasal instillation (IN).

**Fig. S6.**
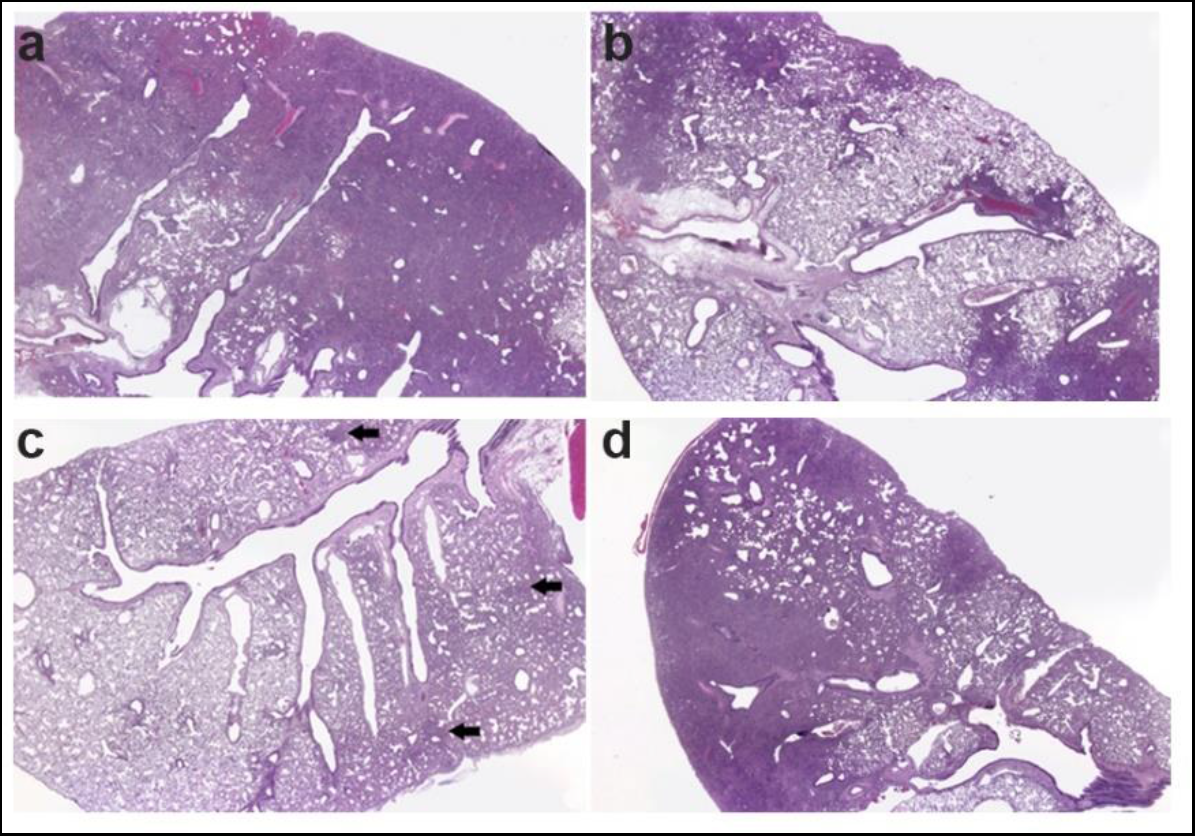
Representative Lung Histology images. Images of a representative hamster lung per group of animals that received a multi-dose of 25 mg/kg of **a)** Ctl siRNA or **b)** COV-siRNA1+2 at 7 and 14 days prior infection or a single dose of COV-siRNA1+2 at 7 days prior infection at c) 25 mg/kg or d) 50 mg/kg. All animals treated with COV-siRNA1+2, had decreased in severity and/or incidence in all SARS-CoV-2-related microscopic findings, except for pleural fibrosis, compared to Ctl siRNA-treated animals. Microscopic findings are concentrated in basophilic (dark blue/purple) consolidated areas, fewer areas were observed in animal c and are depicted by arrows

**Table S1.**
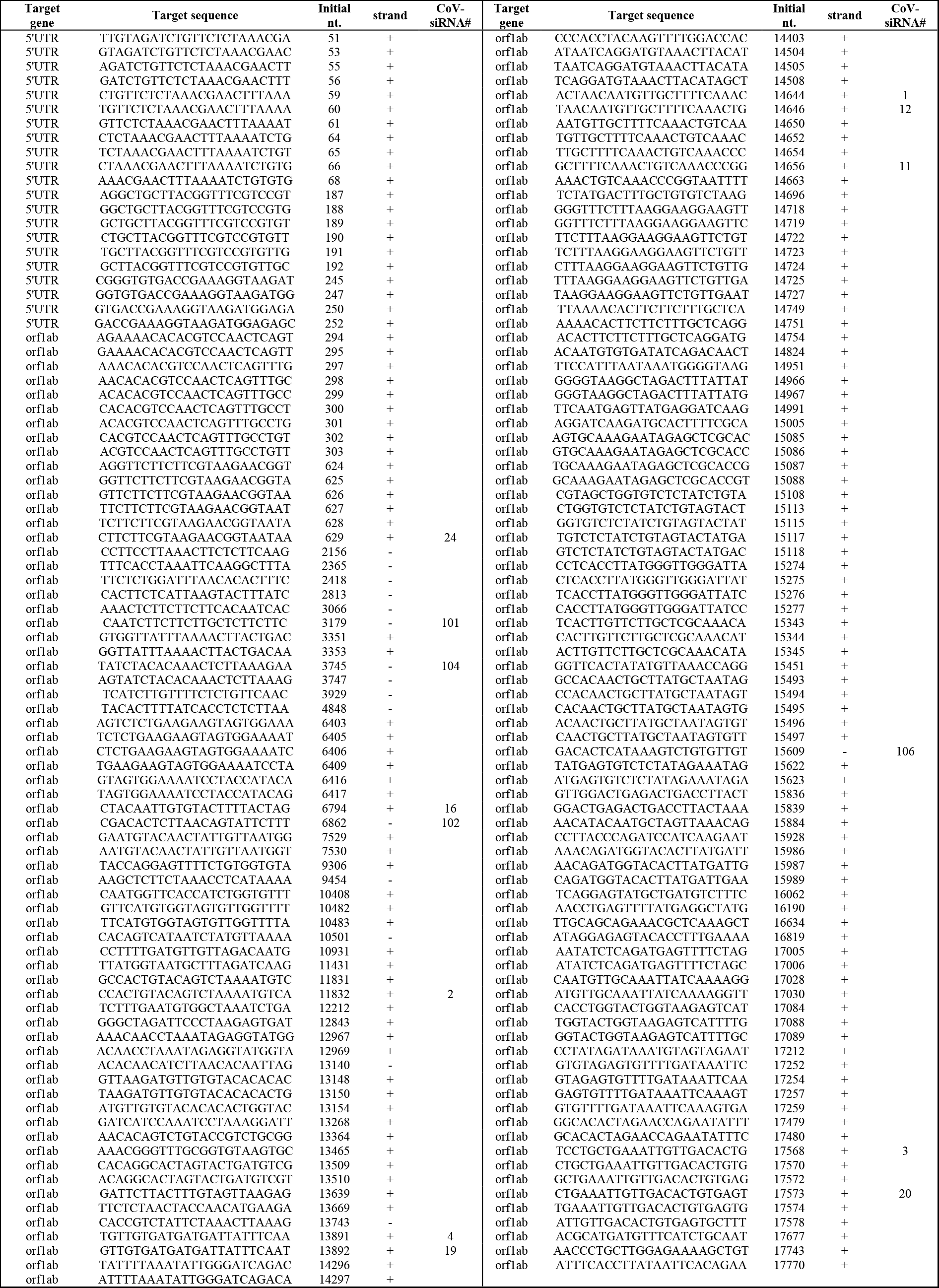
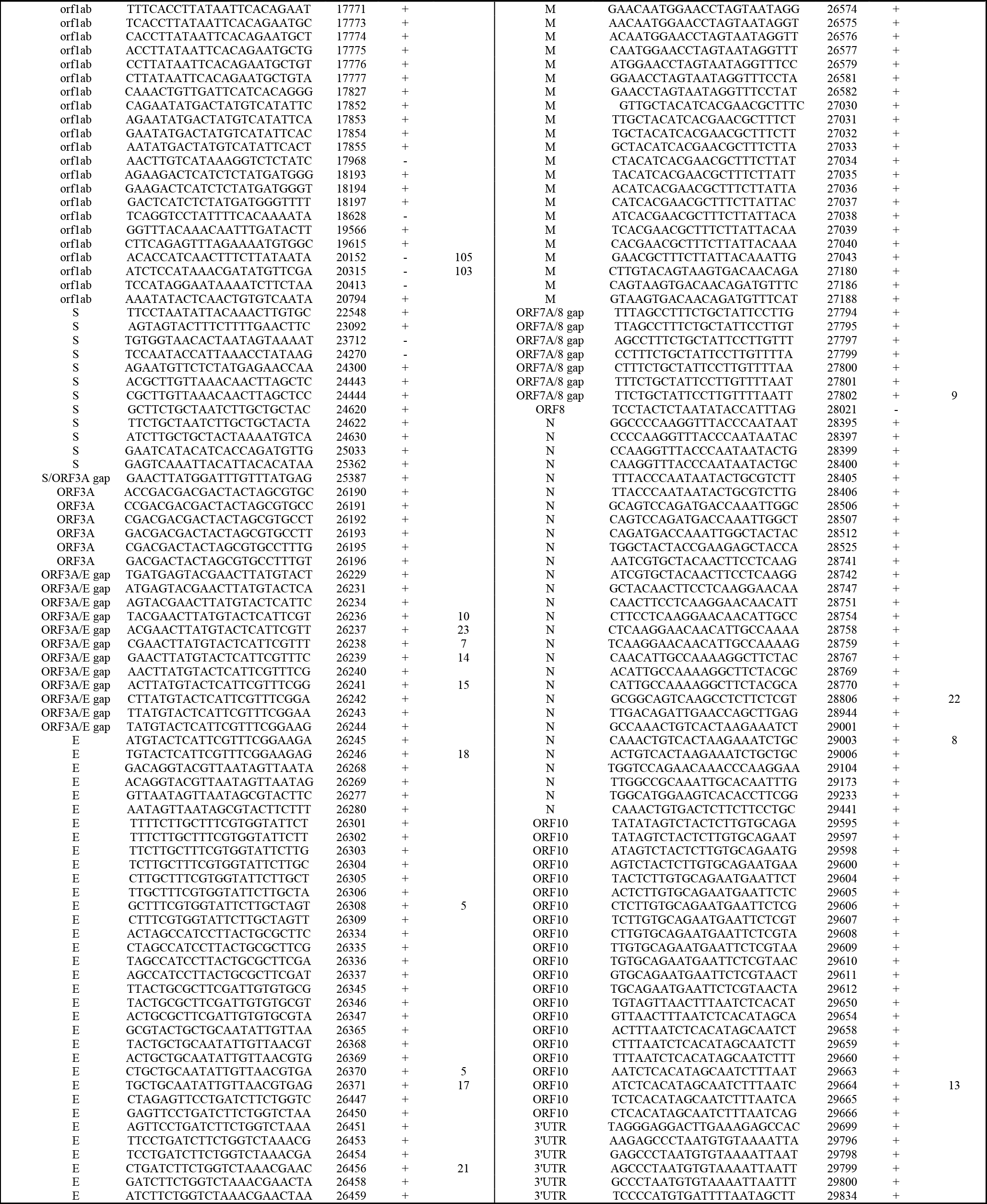
siRNAs target sequences.

**Table S2.**
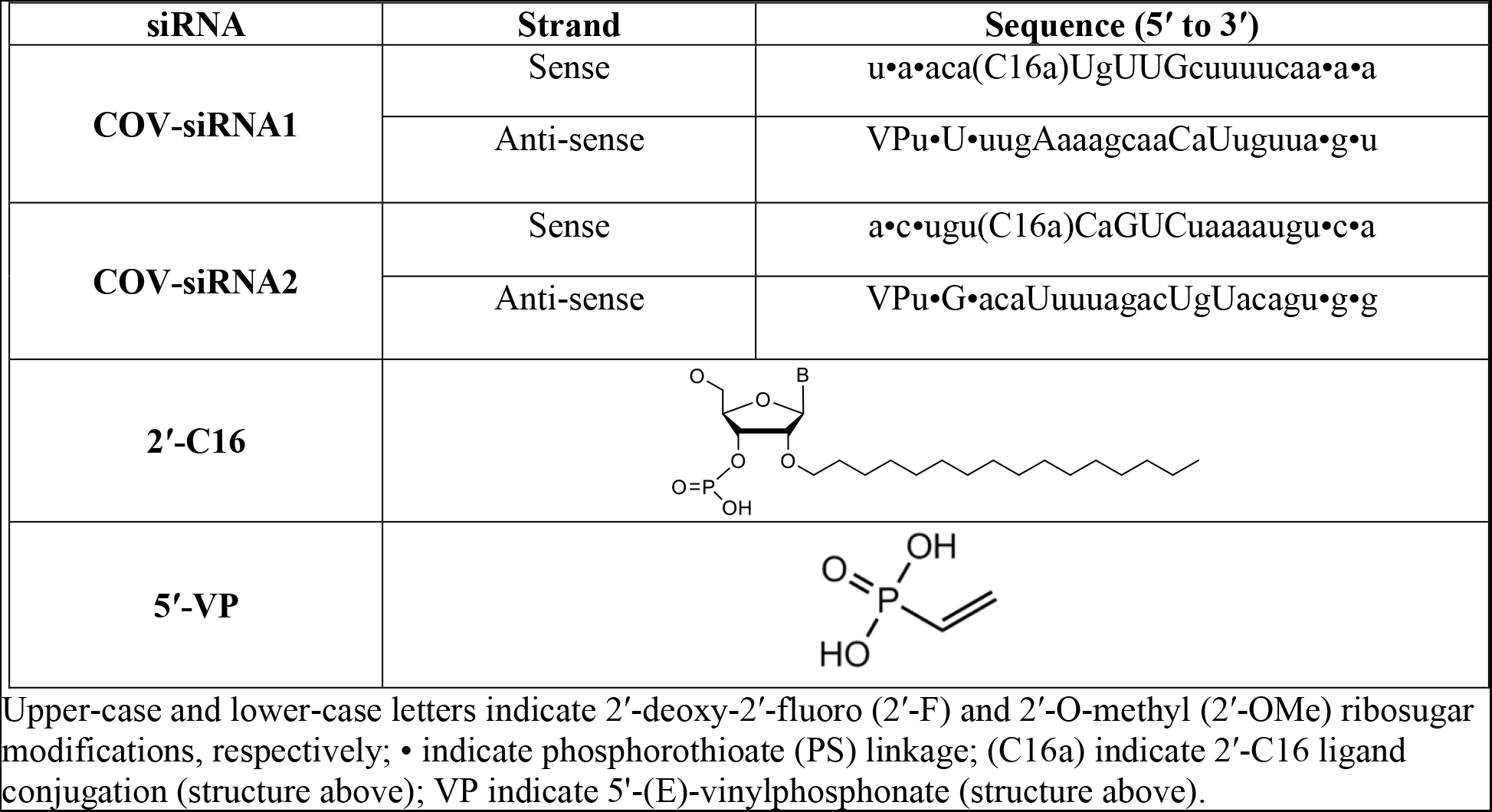
Designs and sequence of the two-siRNAs used in combination

**Table S3.**
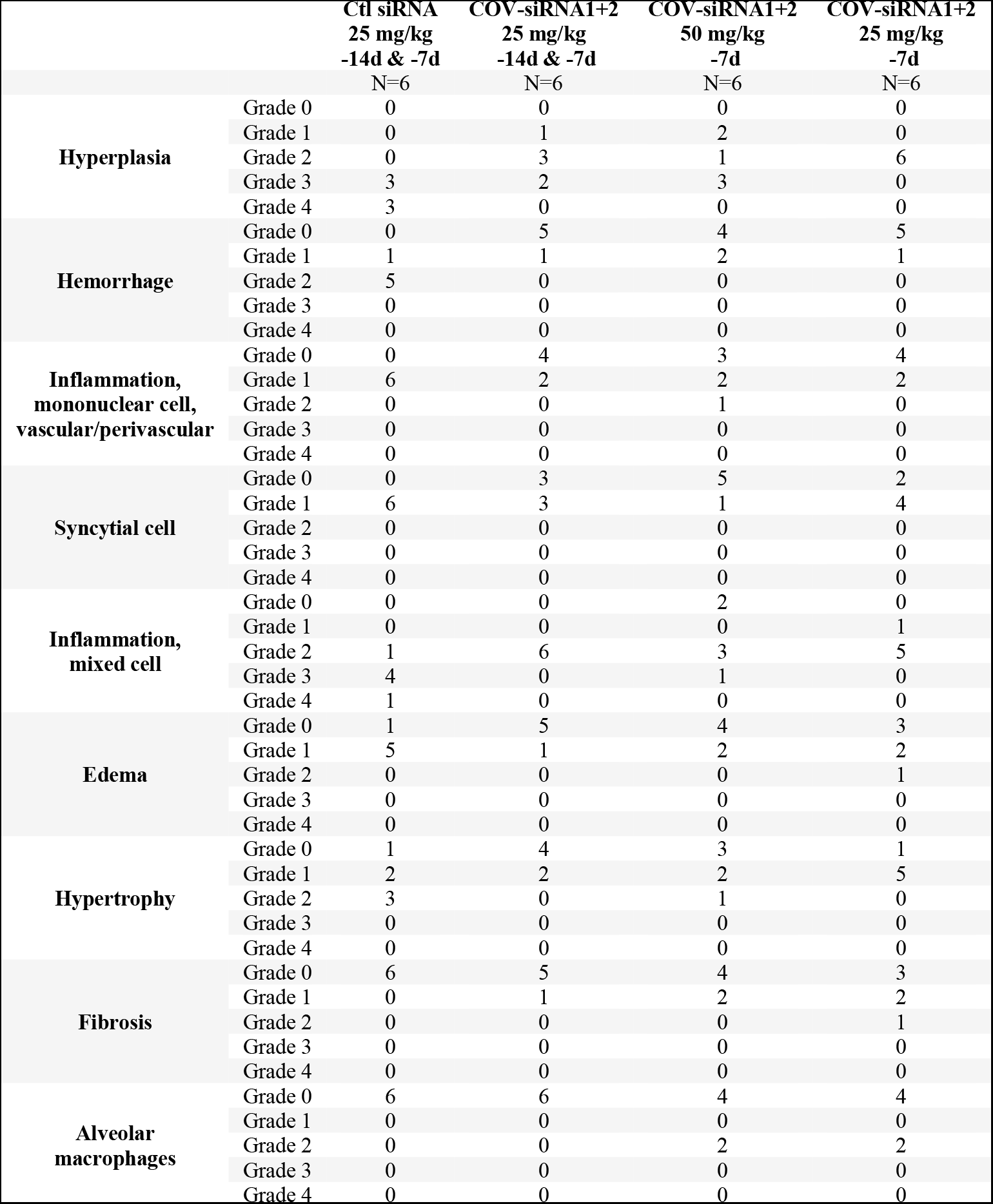
Incidence summary of hamster lung microscopic findings

